# Evaluating the Fidelity of Data-Driven Predator-Prey Models: A Dynamical Systems Analysis

**DOI:** 10.1101/2024.11.15.623759

**Authors:** Anna-Simone J. Frank, Jiawen Zhang, Sam Subbey

**Affiliations:** Computational Biology Unit, Department of Informatics, University of Bergen, Bergen, Norway; School of Computation, Information and Technology, Technical University of Munich, Garching, Germany; Institute of Marine Research, Bergen, Norway

**Keywords:** Predator-prey dynamics, SINDy algorithm, Bifurcation analysis, Sensitivity analysis, Ecological systems, Model validation

## Abstract

In empirical predator-prey systems, understanding the inherent dynamics typically comes from analyzing a structural model fitted to observation data. However, determining an appropriate model structure and its parameters is often complex and highly uncertain. A promising alternative is to learn the model structure directly from time series data of both predator and prey.

This study explores the capability of a data-driven algorithm, Sparse Identification of Nonlinear Dynamics (SINDy), to accurately capture the dynamics of a predator-prey system. We apply SINDy to derive a Learned Model (LM) from data generated by a Reference Model (RM), whose predator-prey dynamics are well understood. The study compares the dynamics of the LM to the RM using criteria such as equilibrium points, stability, sensitivity, and bifurcation analysis.

Our results demonstrate general consistency between the RM and LM dynamics, though notable differences remain. We discuss the implications of these differences in the broader context of using learned models to uncover the inherent drivers of predator-prey dynamics and ecological implications.

**2010 MSC:** 37M05, 37N25, 92B05, 92D25, 92D40, 65L05, 37G15

**Highlights:**

- Evaluates data-driven model’s ability to replication of predator-prey dynamics using SINDy framework
- Learned Model captures core dynamics but shows parameter sensitivity and bifurcation differences
- Analysis reveals need for extensive data for effective model learning
- Suggests combining data-driven methods with biological priors to improve model accuracy

## 1. Introduction

Understanding the dynamics of species interactions in empirical marine ecosystems is non-trivial. This is because the interactions occur on different non-deterministic species aggregation and space-time scales[1, 2, 3, 4]. By their nature, it is challenging to define models that adequately capture the inherent dynamics of marine ecosystems [5, 6]. Food webs (FW) are descriptive diagrams of biological communities within an ecosystem, with focus on interactions between predators and prey (or consumers and resources [7]). They can be considered as idealized representations of ecosystem complexity that captures species interactions and community structure, as well as the inherent processes and drivers that determine the dynamics of energy transfer in the ecosystem [8]. If correctly defined, food web models (FWM), such as, Lotka-Volterra models [9], can provide information about the inherent predator-prey process dynamics. Such information is derived using a combination of tools including (but not limited to) bifurcation analysis and stability of equilibrium points [10].

While mathematical models are useful in understanding the system dynamics [1], limited knowledge and partially inaccurate assumptions [11], can lead to structural model uncertainties. In predator-prey models, structural uncertainty might arise if the model does not capture certain biological mechanisms (e.g., time lags in responses to population changes, density-dependent effects, or environmental variability). Several models may be proposed to represent these dynamics, but structural uncertainty persists if there is no clear evidence about which one is correct. Furthermore, even a well-calibrated model with accurate parameter estimates may still give inaccurate predictions if the chosen model structure is not reflective of the actual system dynamics.

Evaluating model adequacy requires a combination of various statistical analyses along with a thorough understanding of the original assumptions and objectives for which the mathematical model was designed and developed [12]. Assessing model adequacy in ecological models is particularly challenging due to the complex, non-linear dynamics of ecosystems, short time series data, and the potential for overfitting, which can obscure the underlying processes. Short time series limit parameter estimation, making it difficult to capture essential trends and leading to uncertainty in predictions. Additionally, measurement errors, variability, and context-dependent interactions complicate model selection and validation.

These challenges discussed above necessitate innovative approaches to model definition. One viable approach is to exploit recent advances in data-driven modeling techniques[13, 14] in learning models from observations. One specific example approach is the Sparse Identification of Nonlinear Dynamics (SINDy)[15]. SINDy is a data-driven algorithm designed to discover governing equations for dynamical systems from time series data. It is based on sparse regression and selects model terms from a candidate library to describe the data pattern. By balancing complexity and accuracy, the risk of overfitting to the data set is reduced [16, 17, 18]. This attribute makes SINDy a preferrable choice compared to methods for deriving the structure of nonlinear dynamical systems from data [13]. However, while SINDy is powerful and widely used, other similar methods exist, such as, Eureqa[14], which is based on symbolic regression. Eureqa searches for the simplest mathematical expressions that best fit the data, offering interpretability similar to SINDy but with the ability to discover novel functional forms. Our choice is based on the fact that SINDy may be considered one of the leading methods for discovering governing equations from data, with application in various fields including physics (modeling physical systems and understanding complex phenomena), biology (in analyzing ecological dynamics and population models) and engineering (designing control systems and predicting system behavior). Thus our motivation to use SINDy as a tool for discovering governing equations from data stems from it broad application.

In predator-prey systems, the ability of a learned model to fit observational data and make short-term predictions is insufficient for understanding the full dynamics of the system. This is because ecological systems are driven by complex, long-term interactions that go beyond immediate trends. Accurate characterization of the inherent dynamics, such as equilibrium points, stability, and bifurcations, is essential for capturing key ecological processes like population cycles, extinction risks, and responses to environmental changes.

Equally important is the need to ensure that a learned model does not introduce dynamics or characteristics that are not inherent to the actual predator-prey system. Such models may over-complicate the system, misrepresent natural behaviors, or suggest the presence of feedback loops, equilibrium points, or thresholds that do not exist in reality. For instance, falsely predicting instability or dramatic population cycles could trigger unnecessary interventions, such as over-harvesting predators or prey. Ensuring that the model reflects only the true dynamics of the system is crucial to avoid misinterpretation. Such misinterpretation can lead to actions (e.g., harvest decisions) that may potentially, but inadvertently, harm the ecological system.

Given these considerations, the objective of this paper is to evaluate how well a learned predator-prey model preserves the inherent dynamics of the system that generated the observational data. Additionally, we examine potential artifacts introduced by the learned model, such as acquiring dynamical properties that do not correspond to the original system. We use a well-established predator-prey model from the literature, represented by a set of coupled Ordinary Differential Equations (ODEs), as our Reference Model (RM). This RM is considered the true predator-prey system, from which we generate synthetic observations. Since the dynamics of the RM are fully known, we can rigorously assess the performance of the Learned Model (LM) by comparing its ability to replicate the true system dynamics.

The article is organized as follows: Section 2 introduces the Reference Model (RM) and provides an overview of the SINDy modeling framework. This section also discusses our specific modeling procedure, which uses a Python implementation of SINDy. In Section 3, we detail the procedure used to analyze the results from the modeling process. Section 4 presents the findings, which are further explored in Section 5. Finally, the conclusions drawn from this research are summarized in Section 6.

## 2. The reference model and SINDy

This section introduces the reference model, and explains the data generating process. We then describe the SINDy algorithm [19].

### 2.1. The reference model and data generation

Our Reference Model, adapted from [20], is represented by the system of coupled ODEs in (1), where (*a, b, c, d, e, h*) defines a set of model parameters. This system describes the interaction between a predator species, *y* (capelin), and its prey, *x* (zooplankton).

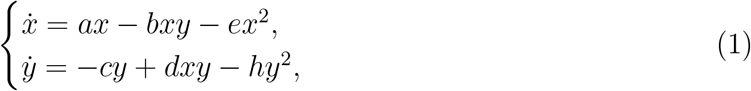

In (1), the term *ax* − *ex*^2^ represents logistic growth of prey with maximum carrying capacity of *a/e*. The parameter *b* indicates the natural mortality rate of the prey *x*. Parameter *c* describes the natural mortality rate of predator *y* and *h* describes its death due to intraspecific competition. Parameter *d* is the growth rate of predator *y* which preys on *x*. The model was calibrated using empirical observations, and the optimized model parameters, taken from [20], are presented in Table 1.

**Table 1:**
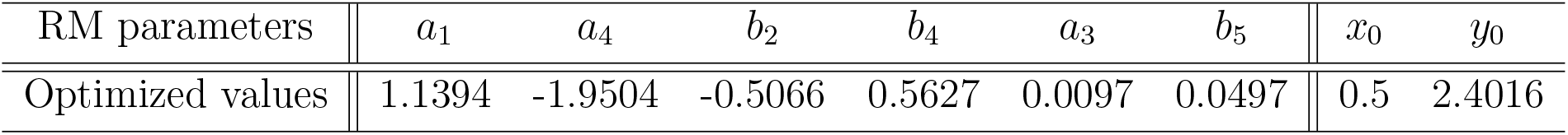
Parameters of the optimized model and initial values (*x*_0_, *y*_0_), taken from [20].

| RM parameters | $a_1$ | $a_4$ | $b_2$ | $b_4$ | $a_3$ | $b_5$ | $x_0$ | $y_0$ |
| --- | --- | --- | --- | --- | --- | --- | --- | --- |
| Optimized values | 1.1394 | -1.9504 | -0.5066 | 0.5627 | 0.0097 | 0.0497 | 0.5 | 2.4016 |

We combine the basis functions for 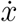 and 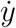, respectively, [1, *x, y, xy, x*^2^] and [1, *y, xy, y*^2^], to derive new expressions for the RM, given by (2). This general structure, allows us to more easily compare the basis functions, across all models.

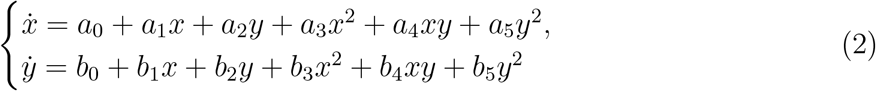

Table 1 gives values of fnon-zero parameters for the generalized model expressions.

To generate our reference time series, we solved the ODE system using the `solve_ivp` function from Python’s ODE solver package (version 2.7.16) [21]. The system was solved over *t* ∈ [0, 40] with a time step of Δ*t* = 0.05.

### 2.2 SINDy– A brief overview

The overview presented in this section is adapted from [15]. Given a set of state variables *x*(*t*) ∈ **R**^*n*^ measured at time points *t*_1_, *t*_2_, …, *t*_*m*_, the SINDy algorithm seeks to reconstruct the governing equations of the system:

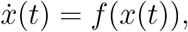

where *f* (*x*) is a sparse nonlinear function describing the system’s dynamics. An essential concept behind SINDy is that the function *f* (*x*) can often be represented as a combination of only a few active terms from a larger set of candidate functions, making it sparse in nature [22].

To determine *f* (*x*), we first approximate the time derivative 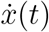 from the measurement data *x*(*t*). We then arrange the data and its derivatives into two matrices:

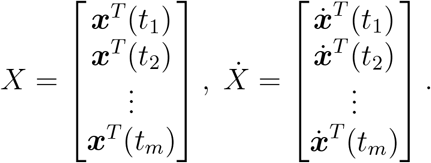

Next, we construct a candidate library of nonlinear functions Θ(*x*), which may include polynomials, trigonometric functions, and other potential system dynamics. The goal is to find the sparse vector of coefficients 𝒞= [*ξ*_1_, *ξ*_2_, …, *ξ*_*n*_] that identifies the active terms in the library, allowing us to express the system’s dynamics as:

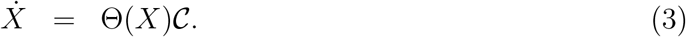

This optimization problem is solved using a sparsity-promoting algorithm, such as least-squares regression with sparsity regularization [16, 17].

To enhance robustness to data noise and improve nonlinear parameter estimation, we employ the Sparse Relaxed Regularized Regression (SR3) algorithm in determining 𝒞, by solving the following optimization problem:

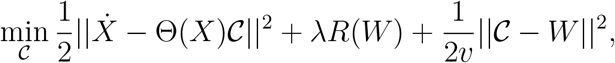

where *W* is an auxiliary variable, *v* is a regularization strength parameter, and *R*(*W*) is a sparsity-inducing regularization term.

Once 𝒞 is determined, the learned governing equations can be written as:

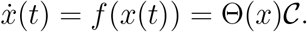

For our analysis, we used the Python implementation of SINDy, provided by the PySINDy package (version 1.7.2) [23, 24].

### 2.3 Algorithm Workflow: Model Selection and Hyperparameter Tuning

The approach adopted is divided into two parts. Part 1 outlines the steps to identify the best governing model equations (LM) from the data. This step focuses on selecting the right structure for the model, specifically by identifying the basis functions or active terms from a library of candidate functions. These basis functions capture the essential relationships in the system, such as interactions between species.

After the model structure is set, Part 2 refines the model (resulting in the Re-estimated Parameter Model (RPM)), by re-estimating the coefficients of the identified basis functions using the data. Since the SINDy algorithm uses derivatives of the data **x**(*t*), rather than the data itself, there may be variability or noise in the calculated derivatives, potentially making the estimated model coefficients (𝒞) less reliable. Re-estimating the coefficients directly from the original data in Part 2 compensates for this potential inaccuracy by fine-tuning the coefficients to better ma tch the ob served data patterns. Since the parameter re-estimation in Part 2 is handled via standard nonlinear least-squares, we omit detailed discussion and concentrate on Part 1, which finds the best governing equations (or active terms in the library, Θ).

The algorithm’s input data is generated from the RM and then divided into training and test sets in a 0.65:0.35 ratio. This ratio, denoted as *r*, corresponds to the point of intersection between the curves representing the RMSE values for model fit to the data and model predictions, both of which are functions of the training data ratio. The RMSE for model fit decreases as the training set ratio increases, while the RMSE for model predictions exhibits the opposite trend. Therefore, this point of intersection can be interpreted as a trade-off between model bias and accuracy [25].

In order to determine the governing model equations, the training set is further divided into training and validation sets. Thus, the data is split into three sets: training, validation, and testing. We apply 3-fold validation on these sets to select optimal hyperparameters in PySINDy [23, 24]. The chosen hyperparameters are: (1) *D*, the numerical differentiation method that approximates 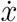 from data *x*; and (2) *T*, the threshold in the differentiation method to control regularization strength. The set of differentiation methods *D* includes `FiniteDifference, finite_difference, SmoothedFiniteDifference, savitzky_golay, spline, trend_filtered`, and `spectral`, while the threshold *T* ranges from 0 to 1 in steps of 0.1. We use a feature library with six second-order polynomial terms: 1, *x, y, x*^2^, *y*^2^, *xy*, where *x* and *y* denote prey and predator, respectively. We employ Sparse Relaxed Regularized Regression (SR3) to solve (3) for 𝒞, under the constraint that *y* consumes *x*.

Algorithm 1 summarizes the procedure for generating the LM and RPM.

#### Algorithm 1

Model Building Algorithm based on PySINDy [24]

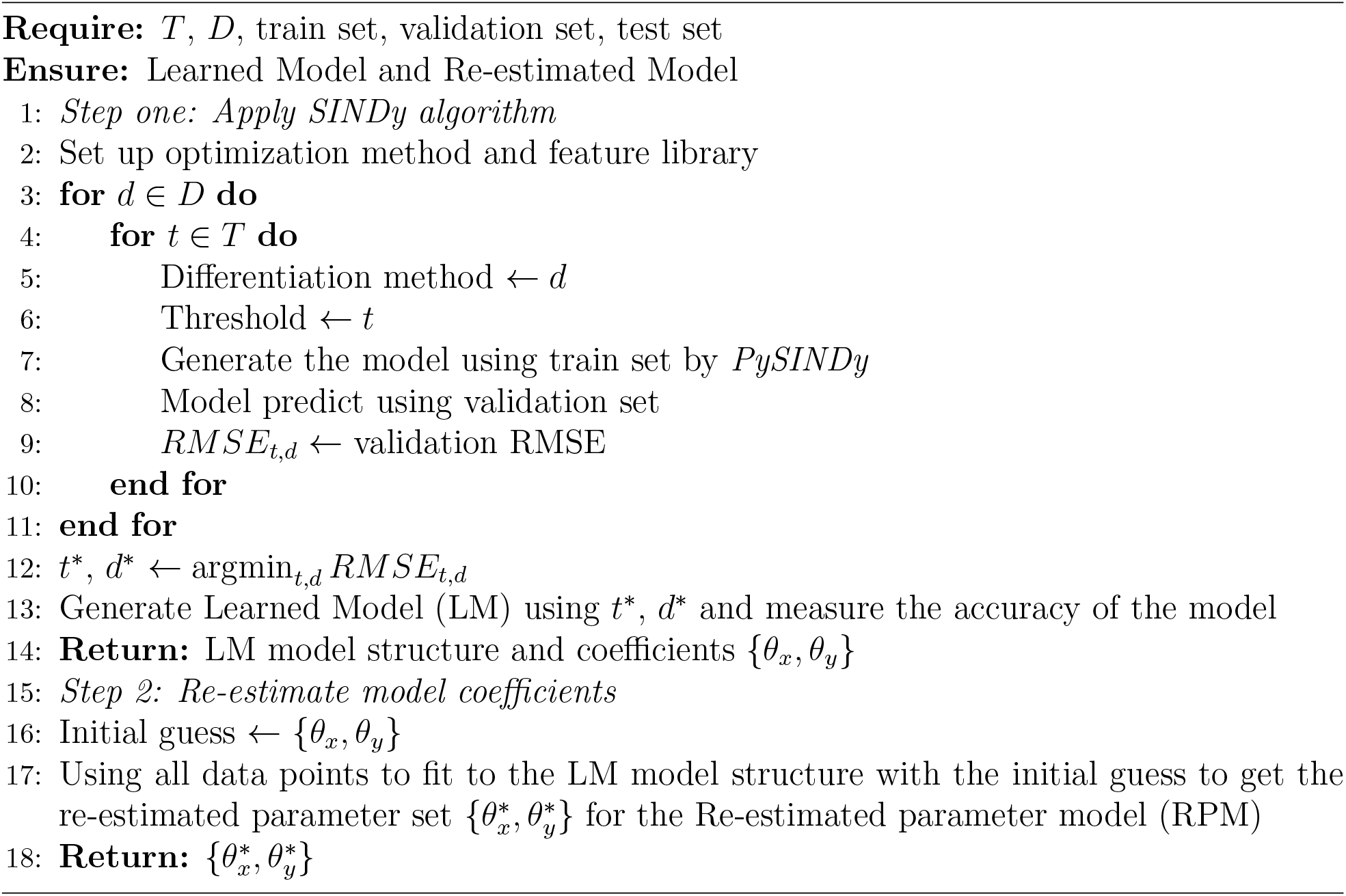

## 3. Model Comparison and Analysis

We perform stability, sensitivity, and bifurcation analysis for all three (RM, LM, RPM) models. The results from these analyses form the basis for comparing and contrasting the models.

### Stability of Equilibrium Points

We first calculate the equilibrium points of the three models, and determine the stability of each equilibrium point under perturbation [26]. While the equilibrium point calculation of the RM is trivial, the other models involved solving for the roots of a quartic equations, which we do using a Python implementation of the method by Ferrari [27]. We used the Routh–Hurwitz stability criterion to determine stability of the equilibrium points [28, 29, 30].

### Model Parameter Sensitivity

We perform sensitivity analysis to assess how model output uncertainty is impacted by various input parameters [31], comparing the LM and RPM models to the RM to determine if they share similarly sensitive parameters. Using Sobol’s method [32, 33, 34] – a variance decomposition approach that handles nonlinear models and parameter interactions – we calculate sensitivity indices with Python’s SALib package (version 1.4.8). Unlike in [20] which focused only on the parameters *a*_3_ and *b*_5_, we examine the impact of all non-zero parameters, adjusting each by ±50%. Using Saltelli sampling, we generate parameter samples (14000, 24000 and 22000 parameter samples) for each model and calculate sensitivity indices across time points, visualizing results with a heat map. Blue squares indicate high sensitivity (index >0.5), while red squares indicate low sensitivity. Finally, we average the sensitivity indices over time to identify the most influential parameters.

### Bifurcation analyses

To compare the qualitative behavior of all three models, we perform single- and two-parameter bifurcation analyses using Python’s PyDStools package (version 0.91.0). Building on the Hopf-bifurcation analysis of parameters *a*_3_ and *b*_5_ in the RM [20], we examine corresponding parameters in the LM and RPM models.

A Hopf-bifurcation describes the emergence of periodic orbits as an equilibrium point’s stability changes, marked by a pair of imaginary eigenvalues [35]. In 2D ODE models, this occurs when the Jacobian trace is zero, according to the Routh–Hurwitz criterion. While the primary focus is Hopf-bifurcation, we also briefly discuss other types that may emerge in single- and two-parameter analyses [36]:

- Saddle-node (also called Limited Point) bifurcation occurs when two equilibria collide and vanish, characterized by a zero eigenvalue.
- Generalized Hopf-bifurcation (GH) occurs when purely imaginary eigenvalues pair with a vanishing first Lyapunov coefficient in a two-parameter ODE model.
- Bogdanov-Takens (BT) bifurcation arises at an equilibrium with a zero eigenvalue of multiplicity two, indicating two colliding equilibrium points.
- Zero-Hopf (ZH) bifurcation occurs at an equilibrium with a zero and a pair of imaginary eigenvalues, marking the intersection of saddle-node and Hopf-bifurcation curves.

## 4. Results

### 4.1. Comparing model structures

The model dynamics and structural differences depend on the specific values of the model parameters, which are detailed in Table 2 for the RM, LM, and PRM.

**Table 2:** Parameter values of (2) for the RM, LM, and RPM.

| Coefficients in $\dot{x}$: | $a_0$ | $a_1$ | $a_2$ | $a_3$ | $a_4$ | $a_5$ |
| --- | --- | --- | --- | --- | --- | --- |
| RM | 0 | 1.1394 | 0 | 0.0097 | -1.9504 | 0 |
| LM | 0.001 | 1.134 | 0.006 | 0.011 | -1.948 | -0.010 |
| RPM | 0.003 | 1.133 | -0.008 | 0.013 | -1.958 | 0.006 |
| Coefficients in $\dot{y}$: | $b_0$ | $b_1$ | $b_2$ | $b_3$ | $b_4$ | $b_5$ |
| RM | 0 | 0 | -0.5066 | 0 | 0.5627 | 0.0497 |
| LM | 0 | 0.003 | -0.569 | -0.001 | 0.562 | 0.053 |
| RPM | 0 | 0.002 | -0.574 | 0 | 0.559 | 0.058 |

From Table 2, we observe that the model coefficients of all three models are similar up to the second decimal place. The LM and RPM models include more terms, such as an intercept (*a*_0_), which the RM model lacks. Overall, the model structures are comparable, particularly considering that the additional terms in the LM and RPM have values close to zero. This similarity is further illustrated in the simulated dynamics of the LM and RPM models compared to the RM model (see Figure 1).

**Figure 1:**
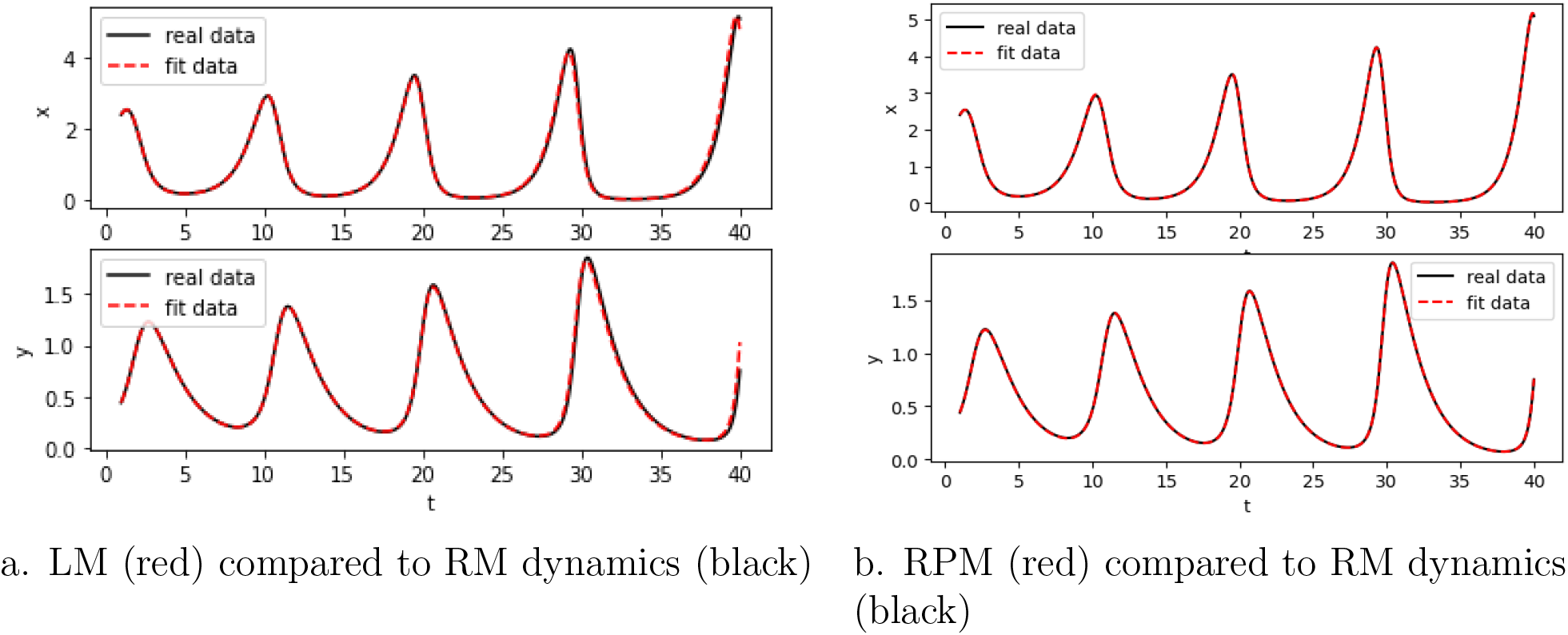
Simulated model dynamics for prey (*x*) and predator (*y*) of the LM, RPM, and RM.

### 4.2 Dynamical system characteristics

As shown in Section 4.1, the model structures and simulations are similar between the RM and the learned equations in the LM and RPM. However, to determine if the data-driven algorithm accurately captures the inherent dynamics of the RM, we will present and compare the dynamical characteristics of all three models.

### Equilibrium points and their stability

Table 3 summarizes the equilibrium points and their stability. The RM has a unique positive equilibrium point, while the LM and RPM have four equilibrium points due to the quartic polynomial. In our comparison, we focus only on the positive equilibrium points, as biomasses cannot be negative.

**Table 3:** Classification of type and stability of coexistent equilibrium points for the RM, LM, and RPM.

| Model structure | Coexisting equilibrium points $(x^*, y^*)$ | Stability and type |
| --- | --- | --- |
| RM | (0.955, 0.589) | unstable spiral |
| LM | (0.951, 0.588) | unstable spiral |
| RPM | (0.960, 0.585) | unstable spiral |
|  | (0.027, 9.634) | unstable saddle node |

Table 3 indicates that both the RM and the LM have a single (coexisting, unstable spiral) equilibrium point located at approximately (0.95, 0.59) when rounded to the second decimal place. In contrast, the RPM features two coexisting equilibrium points: (i) an unstable spiral at approximately (0.96, 0.59), and (ii) an unstable saddle node at approximately (0.03, 9.63), also rounded to the second decimal place. The equilibrium point at around (*x*^*^, *y*^*^) = (0.95,0.59) is the same for all three models.

### Model parameter sensitivity

Table 4 presents the most sensitive parameters of the three models, ranked in descending order based on the mean total effect of Sobol’s sensitivity index. The sensitivity parameters for each model are also included in the table. In the RM and LM, the sensitivity parameters are identical, with similar index values. However, in the RPM, parameter *b*_5_ (corresponding to *y*^2^ in 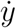) is sensitive, while in the RM and LM, it is *b*_2_ (corresponding to *y* in 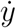).

**Table 4:**
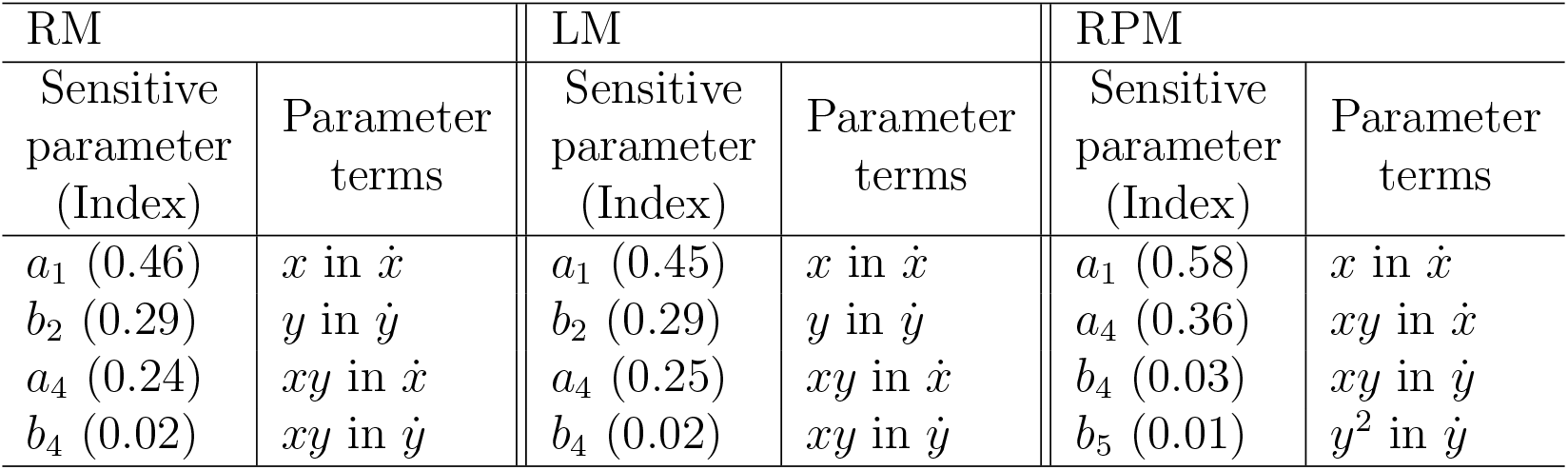
The four most sensitive model parameters in descending order for the RM, LM, and RPM. “Index” indicates the mean of Sobol’s sensitivity index over the time series, reflecting the total effect.

### Bifurcations in model dynamics

#### Single-parameter Hopf-bifurcations in all three models

In the analysis of the RM, the Hopf-bifurcation occurs for the parameter *a*_3_ of term *x*^2^ in 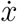 equation and parameter *b*_5_ of the term *y*^2^ in 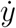 equation.

Similar to the analysis of the RM model in (1), we analyzed also the other two models for the occurrence of Hopf-bifurcation around the positive coexisting equilibrium point (*x*^*^, *y*^*^) = (0.95, 0.59) with respect to the four most sensitive model parameters (*a*_1_, *b*_2_, *a*_4_, *b*_4_), as well as the Hopf-bifurcation parameters (*a*_3_, *b*_5_) from [20]. In all three models, Hopf-bifurcation occurs for the parameters *a*_3_ and *b*_5_. Table 5 shows the values of critical bifurcation points for *a*_3_ and *b*_5_, which are similar when rounded to the second digit point. For all three models, it can be observed that crossing the critical Hopf-bifurcation point changes stability of the equilibrium point (*x*^*^, *y*^*^) = (0.95, 0.59) from unstable to stable.

**Table 5:** Table of critical Hopf-bifurcation points for all models.

| Model/Parameters | $a_3$ | $b_5$ |
| --- | --- | --- |
| RM | -0.0296 | -0.0169 |
| LM | -0.027 | -0.0114 |
| RPM | -0.03 | -0.01637 |

#### Comparison of two-parameter bifurcations

Only parameter *a*_1_ shows Hopf-bifurcation dynamics in both LM and RPM (Tables 6–8). All other parameters not appearing in the RM, specifically *a*_0_, *a*_2_, *a*_5_, *b*_1_, *b*_3_, exhibit single-parameter Hopf-bifurcations (H) with limit points (LP) for *a*_0_ and *b*_3_. An ‘X’ denotes no bifurcation.

**Table 6:**
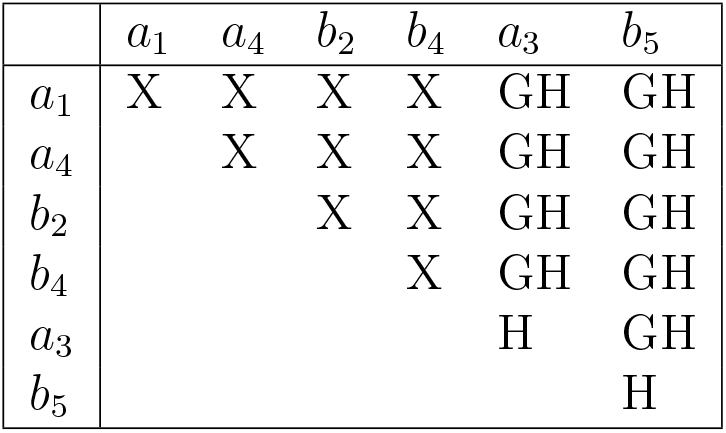
Single and two-parameter bifurcation analysis for RM, with parameters in brackets representing those from (1) (sorted accordingly). “H” denotes a Hopf-bifurcation, “GH” a Generalized Hopf-bifurcation, and “X” indicates the absence of a bifurcation.

| | $a_1$ | $a_4$ | $b_2$ | $b_4$ | $a_3$ | $b_5$ |
| --- | --- | --- | --- | --- | --- | --- |
| $a_1$ | X | X | X | X | GH | GH |
| $a_4$ | | X | X | X | GH | GH |
| $b_2$ | | | X | X | GH | GH |
| $b_4$ | | | | X | GH | GH |
| $a_3$ | | | | | H | GH |
| $b_5$ | | | | | | H |

Off-diagonal elements in Tables 6–8 show two-parameter bifurcations. For the RM (Table 6), Generalized Hopf-bifurcations (GH) occur between *a*_3_, *b*_5_, and other parameters. In contrast, LM (Table 7) shows both GH and Bogdanov-Takens (BT) bifurcations across various parameter combinations.

**Table 7:** Single and two-parameter bifurcation analysis for the LM. “H” denotes a Hopf-bifurcation, “GH” a Generalized Hopf-bifurcation, “BT” a Bogdanov-Takens bifurcation, “LP” a limit point, and “X” indicates no bifurcation.

| | $a_0$ | $a_1$ | $a_2$ | $a_3$ | $a_4$ | $a_5$ | $b_1$ | $b_2$ | $b_3$ | $b_4$ | $b_5$ |
| --- | --- | --- | --- | --- | --- | --- | --- | --- | --- | --- | --- |
| $a_0$ | H, LP | X | BT | BT | X | GH | GH | X | GH,BT | X | X |
| $a_1$ | | H | X | GH | BT | X | BT | BT | BT | BT | GH |
| $a_2$ | | | H | X | X | GH | BT | X | GH,BT | X | X |
| $a_3$ | | | | H | GH | X | BT | X | BT | X | X |
| $a_4$ | | | | | X | X | X | X | X | X | X |
| $a_5$ | | | | | | H | BT | X | GH,BT | X | X |
| $b_1$ | | | | | | | H | X | GH,BT | X | X |
| $b_2$ | | | | | | | | X | X | X | X |
| $b_3$ | | | | | | | | | H,LP | X | X |
| $b_4$ | | | | | | | | | | X | X |
| $b_5$ | | | | | | | | | | | H |

While RPM bifurcations are similar to LM, they differ in occurrence and type, with a Zero-Hopf (ZH) bifurcation noted for (*a*_0_, *a*_3_) (Table 8).

**Table 8:** Single and two-parameter bifurcation analysis for the RPM. “H” denotes a Hopf-bifurcation, “GH” a Generalized Hopf-bifurcation, “BT” a Bogdanov-Takens bifurcation, “ZH” a Zero-Hopf bifurcation, “LP” a limit point, and “X” indicates no bifurcation.

| | $a_0$ | $a_1$ | $a_2$ | $a_3$ | $a_4$ | $a_5$ | $b_1$ | $b_2$ | $b_4$ | $b_5$ |
| --- | --- | --- | --- | --- | --- | --- | --- | --- | --- | --- |
| $a_0$ | H, LP | BT | X | GH,BT,ZH | X | GH | GH,BT | X | X | GH |
| $a_1$ | | H | X | X | BT | X | BT | X | X | X |
| $a_2$ | | | H | GH | X | GH | BT | X | X | GH |
| $a_3$ | | | | H | X | GH | GH,BT | X | X | X |
| $a_4$ | | | | | X | X | X | X | X | X |
| $a_5$ | | | | | | H | BT | X | X | GH |
| $b_1$ | | | | | | | H | X | X | GH |
| $b_2$ | | | | | | | | X | X | X |
| $b_4$ | | | | | | | | | X | X |
| $b_5$ | | | | | | | | | | H |

## 5. Discussion

The main goal of this paper has been to evaluate how well a data-driven model, can replicate the inherent dynamics of a known predator-prey system, using the SINDy modeling framework. By comparing the Learned Model (LM) with a Reference Model (RM) whose dynamics are well-understood, we have assessed the ability of a LM to capture key system dynamic behavior of the system, such as equilibrium points, stability, sensitivity, and bifurcation behaviors. Ultimately, we have sought to determine whether data-driven models can serve as reliable tools for understanding complex ecological dynamics.

Our results show that the LM, generated through the SINDy algorithm, effectively captures the primary dynamics of the RM, including equilibrium points, stability characteristics, and some bifurcation behaviors. This consistency suggests that data-driven approaches like SINDy are promising for replicating known dynamical systems. However, certain discrepancies between the models, particularly in parameter sensitivities and bifurcation types, emphasize the importance of model validation.

Our sensitivity analysis reveals notable variation in the influence of certain parameters on model output. For instance, parameters *a*_1_ and *b*_2_, which are essential in RM, remain influential in LM but display slight differences in index values. This variation indicates that SINDy-based models might shift parameter importance, potentially altering ecological predictions. Such shifts are critical in predator-prey systems, where small changes in growth or mortality rates can significantly impact system stability. Sensitivity to parameter variations highlights that care must be taken in using LMs as a basis for ecological inference.

Results from our bifurcation analysis show that the models exhibit Hopf-bifurcations for some shared parameters, though distinctions emerge with complex bifurcations. For example, the RPM model introduces Zero-Hopf and Bogdanov-Takens bifurcations, which do not appear in the RM. Hence, while SINDy captures primary bifurcation dynamics, some emergent behaviors may not reflect the real predator-prey system. The concern is that the complex bifurcations introduced by the LM can lead to cyclic behaviors or multiple equilibria, which can potentially causing the LM to suggest more unstable dynamics than in the true (reference) system.

The analysis revealed that data on the order of 10^3^ is essential for the LM to effectively capture trends in input data. This translate to thousands of years of ecological observations, which limits the practical application of learning algorithms in this field. A potential solution is to employ compressive sensing (CS)[37], a signal processing technique that enables the recovery of sparse signals using significantly fewer measurements than traditional methods. By enriching the dataset, one could then rely on learning algorithms (e.g., SINDy) to provide the model structure for the dynamics. Once the structure is derived, the model parameters could be re-estimated (similar to our RPM), to obtain parameter values that are consistent with observations. In adopting such a procedure, however, it is crucial to ensure that the data-enrichment process does not introduce biases that could compromise the learning process, resulting in a model that misrepresents the ecological system.

## 6. Conclusions

Data-driven methods like SINDy are valuable for learning dynamics from observational data without needing extensive prior knowledge of system structure. Model-based inference can be misleading however, when a learned model introduces artificial dynamics, such as, bifurcations or exaggerated sensitivity.

Our work underscores the necessity of improving model accuracy, potentially through hybrid approaches that combine data-driven methods with biological priors to minimize artifacts. This necessitates modeling frameworks that balance data-driven learning with ecological realism. Additionally, adapting SINDy to account for environmental variability or stochastic effects could enhance its applicability to ecological systems, which often face unpredictable changes.

## Acknowledgement

This project was part of a SQuaRE at the American Institute of Mathematics (AIM), and the authors thank AIM for its support. Special thanks also to Prof. Christina Kuttler (TUM) for her invaluable feedback.

Jiawen Zhang is grateful for the funding received through the Erasmus+ Programme, which enabled a six-month stay at the Institute of Marine Research (Bergen, Norway), where portions of this work were carried out.

## Data statement

This manuscript uses no biological or experimental data. All figures can be reproduced using the models, the parameters and numerical specifications presented in this manuscript.

## CRediT authorship contribution statement

**Anna S Frank:** Conceptualization, Methodology, Investigation, Writing - original draft, Writing - review & editing, Supervision. **Jiawen Zhang:** Formal analysis, Investigation, Writing - review & editing. **Sam Subbey:** Conceptualization, Methodology, Investigation, Writing - original draft, Writing - review & editing, Supervision.

